# Convergent Acquisition of Glucomannan β-galactosyltransferases in Asterids and Rosids

**DOI:** 10.1101/2024.06.11.597938

**Authors:** Konan Ishida, Matthew Penner, Kenji Fukushima, Yoshihisa Yoshimi, Louis F.L. Wilson, Alberto Echevarría-Poza, Li Yu, Paul Dupree

## Abstract

β-Galactoglucomannan (β-GGM) is a primary cell wall polysaccharide in rosids and asterids. The β-GGM polymer has a backbone of repeating glucose and mannose, usually with mono- or di-galactosyl sidechains on the mannosyl residues. CELLULOSE SYNTHASE-LIKE 2 (CSLA2), MANNAN α-GALACTOSYLTRANSFERASE (MAGT), and MANNAN β-GALACTOSYLTRANSFERASE (MBGT) are required for β-GGM synthesis in *Arabidopsis thaliana*. The single MBGT identified so far, *At*MBGT1, lies in glycosyltransferase family 47A subclade VII, and was identified in Arabidopsis. However, despite the presence of β-GGM, an orthologous gene is absent in tomato (*Solanum lycopersicum*), a model asterid. In this study, we screened candidate *MBGT* genes from the tomato genome, functionally tested the activities of encoded proteins, and identified the tomato MBGT (*Sl*MBGT1) in GT47A-III. Interestingly therefore, *At*MBGT1 and *Sl*MBGT1 are located in different GT47A subclades. Further, phylogenetic and glucomannan structural analysis from different species raised the possibility that various asterids possess conserved MBGTs in GT47A-III, indicating that MBGT activity has been acquired convergently among asterids and rosids. Although functional convergence was observed, the acquired amino acid substitutions among the two MBGT groups were not shared, suggesting different evolutionary pathways to achieve the same biochemical outcome. The present study highlights the promiscuous emergence of donor and acceptor preference in GT47A enzymes, and suggests an adaptive advantage for eudicots to acquire β-GGM β-galactosylation.

## Introduction

Mannans are a class of polysaccharides conserved from algae to eudicots but with large structural variations. The simplest structure, homomannan, consisting of β-(1,4)-linked mannosyl residues (Man), is found as a storage polysaccharide in plants such as *Aloe vera* and ivory nuts, and as a cell wall polysaccharide in some green algae and red algae (Aspinall, 1959; Preston, 1979; Painter, 1983; de O. Petkowicz *et al*., 2001; Moreira and Filho, 2008). Glucomannan, whose backbone is interspersed with β-(1,4)-linked glucose (Glc) and whose mannosyl residues are often acetylated, is also seen as a storage polysaccharide (*e.g.,* konjac, orchid) (Gille and Pauly, 2012; He *et al*., 2017; Voiniciuc, 2022). α-Galactosyl (Gal) substitution on C6 of Man is seen in some storage mannans such as in legume seeds (Meier and Reid, 1982; McCleary *et al*., 1985). Acetylated galactoglucomannan (AcGGM) is widespread in the cell walls of woody tissues of vascular plants, and is particularly abundant in conifers (Capek, Alföldi and Lisková, 2002; Handford *et al*., 2003; Goubet *et al*., 2009; Terrett *et al*., 2019; Chernova *et al*., 2020). Recently, a novel type of galactoglucomannan with a repeating Man-Glc disaccharide unit in the backbone was discovered (Yu *et al*., 2018). The patterned galactoglucomannan can be not only monogalactosylated as found in AcGGM, but di-galactosylated on the Man residues through a β-(1,2)-linkage on α-(1,6)-Gal. This glucomannan is called β-galactoglucomannan (β-GGM) (Yu *et al*., 2022). β-GGM is widespread in eudicots, being found in the asterid and rosid clades, including Arabidopsis, apples, kiwifruit, and tomatoes (*Solanum lycopersicum*) (Schröder *et al*., 2001; Yu *et al*., 2022), but has not been found in monocots (Ishida *et al*., 2023).

Mannan structural variation is likely to be related to physiological function in ways that are only partly understood. In algae, homomannan can serve as a major fibrillar component of the cell wall similar to cellulose in land plants (Preston, 1979; Painter, 1983; Fernández *et al*., 2012). The replacement of Man backbone residues with Glc, and the addition of acetyl groups on the backbone, or branching by α-Gal, is thought to affect the polysaccharide solubility (Scheller and Ulvskov, 2010), and interaction with other cell wall components (Yu *et al*., 2018). It had been thought until recently that the importance of mannans may have decreased in land plant evolution as xyloglucan became more abundant in cell walls (Scheller and Ulvskov, 2010). However, mannans are likely to be essential to some aspects of plant physiology since the putative glucomannan synthase mutant *csla7* shows embryonic lethality and defective pollen tube growth (Goubet *et al*., 2003, 2009). Indeed, challenging the idea of a waning importance of glucomannan through vascular plant evolution is the fact that eudicots acquired the more complex β-GGM during angiosperm evolution (Yu *et al*., 2022; Grieß-Osowski and Voiniciuc, 2023; Ishida *et al*., 2023). The significance of β-GGM seems to be obscured in many tissues by the presence of xyloglucan (Yu et al., 2022). As there is a substantial gap in our understanding of the relationship between mannan structures and physiological functions, it is essential to expand our knowledge about the biosynthesis of β-GGM.

Four types of transferase enzymes are involved in mannan biosynthesis: CELLULOSE SYNTHASE-LIKE A (CSLA, GT2), MANNAN α-GALACTOSYLTRANSFERASE (MAGT, GT34), MANNAN β-GALACTOSYLTRANFERASE (MBGT, GT47), and MANNAN *O*-ACETYLTRANSFERASE (MOAT). Furthermore, the participation of the GT106 putative glycosyltransferase enzymes in mannan production has been identified (Wang *et al*., 2013; Voiniciuc *et al*., 2019) but the activity and function remain unclear. The two major types of mannan are synthesised by different combinations of enzymes in Arabidopsis (Goubet *et al*., 2009; Voiniciuc *et al*., 2015; Yu *et al*., 2018, 2022; Zhong, Cui and Ye, 2019). β-GGM is synthesised by *At*CSLA2, *At*MAGT1, and *At*MBGT1 whereas AcGGM of primary and secondary cell walls is synthesised by *At*CSLA9, *At*MAGT1, and *At*MOAT1-4. Among the four mannan synthesis enzymes, CSLA and MOAT are highly conserved in land plants (Liepman *et al*., 2007; Zhong, Cui and Ye, 2019). However, close orthologues of *At*MAGT1 and *At*MBGT1 are not found in bryophytes and vascular seedless plants (Ishida *et al*., 2023). Consistent with the genomic/phylogenetic information, mannan α-Gal side chains have not been found in *Physcomitrium patens* (Ye and Zhong, 2022).

β-GGM shares the GT families for biosynthesis with xyloglucan (Yu *et al*., 2022; Grieß-Osowski and Voiniciuc, 2023). The β-1,4-glucan backbone is synthesised by CELLULOSE SYNTHASE-LIKE C in the GT2 family and xylosyl side chains are added by xyloglucan xylosyltransferase in the GT34 family. The addition of the second side chains (equivalent to β-1,2-Gal of β-GGM) is catalysed by enzymes in GT47A, which is divided into seven subclades (I–VII) (Yu *et al*., 2022). Interestingly, GT47A has many xyloglucan-modifying enzymes with varying donor preferences but there is no clear correlation between the activity and their position in GT47A. For example, *At*MUR3 and *At*XLT2 xyloglucan galactosyltransferases locate to GT47A-VI and GT47A-III, respectively, albeit both with galactosyltransferase activities. *Sl*XST xyloglucan arabinofuranosyltransferases and *Cc*XBT and *Vc*XBT xyloglucan β-xylosyltransferases are classified in GT47A-III along with *At*XLT2 (Schultink *et al*., 2013; Immelmann *et al*., 2023; Wilson *et al*., 2023). Two highly similar GT47A enzymes from *P. patens* in GT47A-I showed distinctive activities: XLT2 activity and XDT (xyloglucan arabinopyranosyl transferase) activities (Zhu, Dama and Pauly, 2018). The range of activities suggests there is relatively frequent evolutionary emergence of new donor preferences in GT47A enzymes that synthesise xyloglucan side chains.

*At*MBGT1 belongs to GT47A-VII and is the only identified MBGT to date. To investigate conservation of MBGT activity in GT47A-VII in monocots and ANA-grade plants, we previously studied rice (*Oryza sativa*) and Amborella (*Amborella trichopoda*) enzymes (Ishida *et al*., 2023). Neither candidate exhibits MBGT activity, and the rice GT47A-VII enzyme was shown to be a xyloglucan galactosyltransferase. Moreover, mannans containing β-Gal side chains were not found in more than 20 species of monocot and ANA-grade tissues studied (Ishida *et al*., 2023) (Fig. 1). These findings support the notion that the acquisition of MBGT activity and therefore β-GGM containing β-Gal side chains is likely to have occured only in eudicots. However, within eudicots the evolutionary acquisition of MBGT activity and hence β-GGM is not yet well studied.

**Figure 1.**
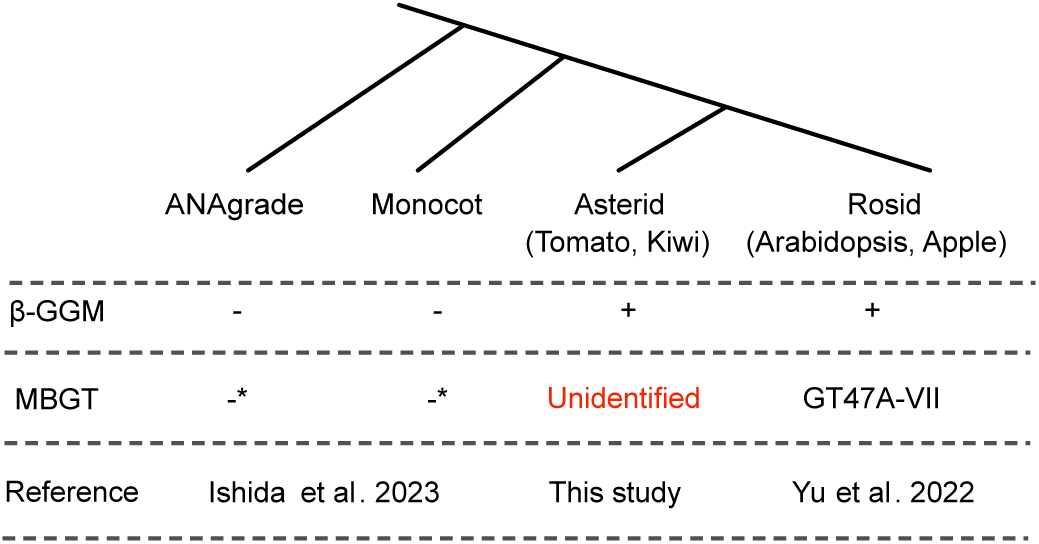
β-GGM and MBGT distribution in the different evolutional stages. (A) Summary of the presence or absence of β-GGM and MBGT enzymes. Asterisks: Genes closest to AtMBGT1 from rice and Amborella did not show MBGT activity *in vivo*.

Here, we searched for an MBGT enzyme from the asterids. Surprisingly, asterids were found to lack a direct orthologue of *At*MBGT1, and the closest homologue in tomato failed to demonstrate MBGT activity. Instead, based on tomato genomic and transcriptomic data, one candidate was found in GT47A-III, a distant and distinct subclade from that of *At*MBGT1. The candidate showed MBGT activity *in vitro* and *in vivo*. Further data support the hypothesis that this MBGT GT47A-III subclade is likely to be conserved throughout asterids. These findings suggest the separate yet convergent acquisition of MBGT in asterids and rosids.

## Results

### Identification of Solyc02g092840 as a candidate asterid MBGT

The MBGT in Arabidopsis is in the GT47A-VII clade (Yu *et al*., 2022). A single GT47A-VII (*Sl*GT11, Solyc03g115750) is found encoded in the tomato genome (Fig. 2A). However, its orthologue in Arabidopsis, *At*GT11, is thought to be a pollen tube-specific xyloglucan galactosyltransferase (Wei *et al*., 2021). Therefore, we re-examined the previous phylogeny (Yu *et al*., 2022) and conducted gene expression analyses to identify any additional MBGT candidates in tomato. Tomato has nine enzymes in the GT47A clade (Fig. 2A). Three of them (*Sl*MUR3 and *Sl*XST1,2) have had their activities confirmed by *in vivo* complementation analysis of the Arabidopsis xyloglucan galactose-lacking mutant *mur3 xlt2* (Schultink *et al*., 2013). The other six are *Sl*GT11, two *At*XUT orthologues in GT47A-II (Solyc08g080930 and Solyc12g056260), one *At*XLT2 orthologue in GT47A-III (Solyc02g092840), one *GT19* orthologue (GT47A-IV; Solyc08g079040), and one *GT17* orthologue (GT47A-V; Solyc08g080930). Interestingly, a previous study reported that *Solyc02g092840* did not complement the xyloglucan galactose-lacking mutant *mur3 xlt2* (Schultink *et al*., 2013), raising the possibility that Solyc02g092840 works on a different substrate. In line with this, gene expression data from fruits showed that *Solyc02g092840* was the most highly expressed GT47A gene through fruit development (that β-GGM is abundant in tomato fruits has been shown in our previous paper (Yu *et al*., 2022)) (Fig. 2B). In contrast, other potential MBGT candidates, such as *SlGT11*, *AtXUT* orthologues, and *SlGT17* had lower expression levels. Hence, we selected Solyc02g092840 as an MBGT candidate.

**Figure 2.**
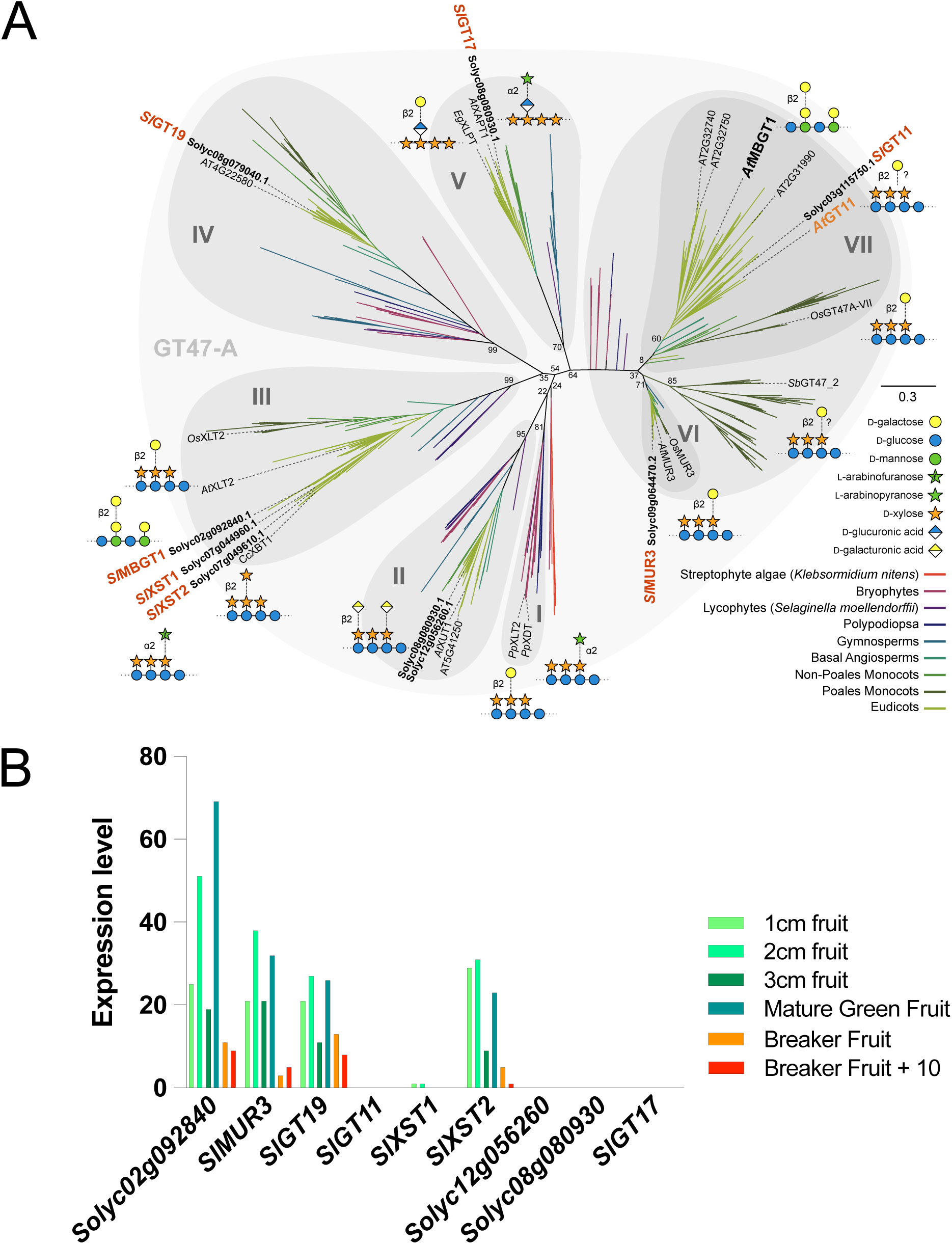
Screening for candidate tomato MBGT. (A) Phylogenetic tree of the GT47A. Relevant GT47A members from tomato are coloured orange. The tree is adapted from our previous publication (Yu *et al*., 2022). MUR3: xyloglucan β-galactosyltransferase. XST: xyloglucan α-arabinofuranosyltransferase. XBT: xyloglucan β-xylosyltransferase. The biochemical activity of *Sl*MUR3, and *Sl*XST1, *Sl*XST2 was confirmed previously (Schultink *et al*., 2013). (B) Tomato GT47A gene expression data in fruits. Original data was obtained from the public database: BAR ePlant tomato (https://bar.utoronto.ca/eplant_tomato/) and are reads per kilobase per million (RPKM) mapped illumina reads.

### Solyc02g092840 exhibits MBGT activity

To investigate β-galactosyltransferase activity of Solyc02g092840 *in vitro* on β-GGM, we transiently expressed the gene *Solyc02g092840* in tobacco (*Nicotiana benthamiana*) leaves. In addition to the candidate, *At*MBGT1 (a positive control) and *Sl*GT11 (the candidate and closest orthologue to *At*MBGT1 in tomato) were also expressed. Microsomal fractions prepared from the transgenic material were subjected to an immunoblot, whereby expression of all three proteins was confirmed (Supplementary Data 1). The microsomal fraction was used as an enzyme source and incubated with UDP-Gal and Arabidopsis seed mucilage containing β-GGM as the substrates (mucilage β-GGM lacks β-Gal because of the trimming by β-galactosidases (Voiniciuc *et al*., 2015, 2016; Yu *et al*., 2018, 2022)). The reaction mixtures were treated with *An*GH5 mannanase for structural characterization of the products. Digestion of the initial substrate yields oligos GA, GM (Fig 3A, where A is Gal-α-(1,6)-Man, G is Glc, and M is Man) (Yu *et al*., 2022). Digestion of *At*MBGT1-treated substrate produced the longer oligos including GBGA and GBGM (where B is Gal-β-(1,2)-Gal-α-(1,6)-Man), because the mannanase cleavage is inhibited at β-galactosylated sites. These oligos were sensitive to β-galactosidase and the main products migrated to GAGA and GAGM (Fig. 3A). Similarly to *At*MBGT1, digestion of Solyc02g092840-treated material also released GBGA and GBGM. In contrast, *Sl*GT11-treated material did not produce β-galactosylated oligos. These results indicate that Solyc02g092840, but not *Sl*GT11, possesses MBGT activity *in vitro*.

**Figure 3.**
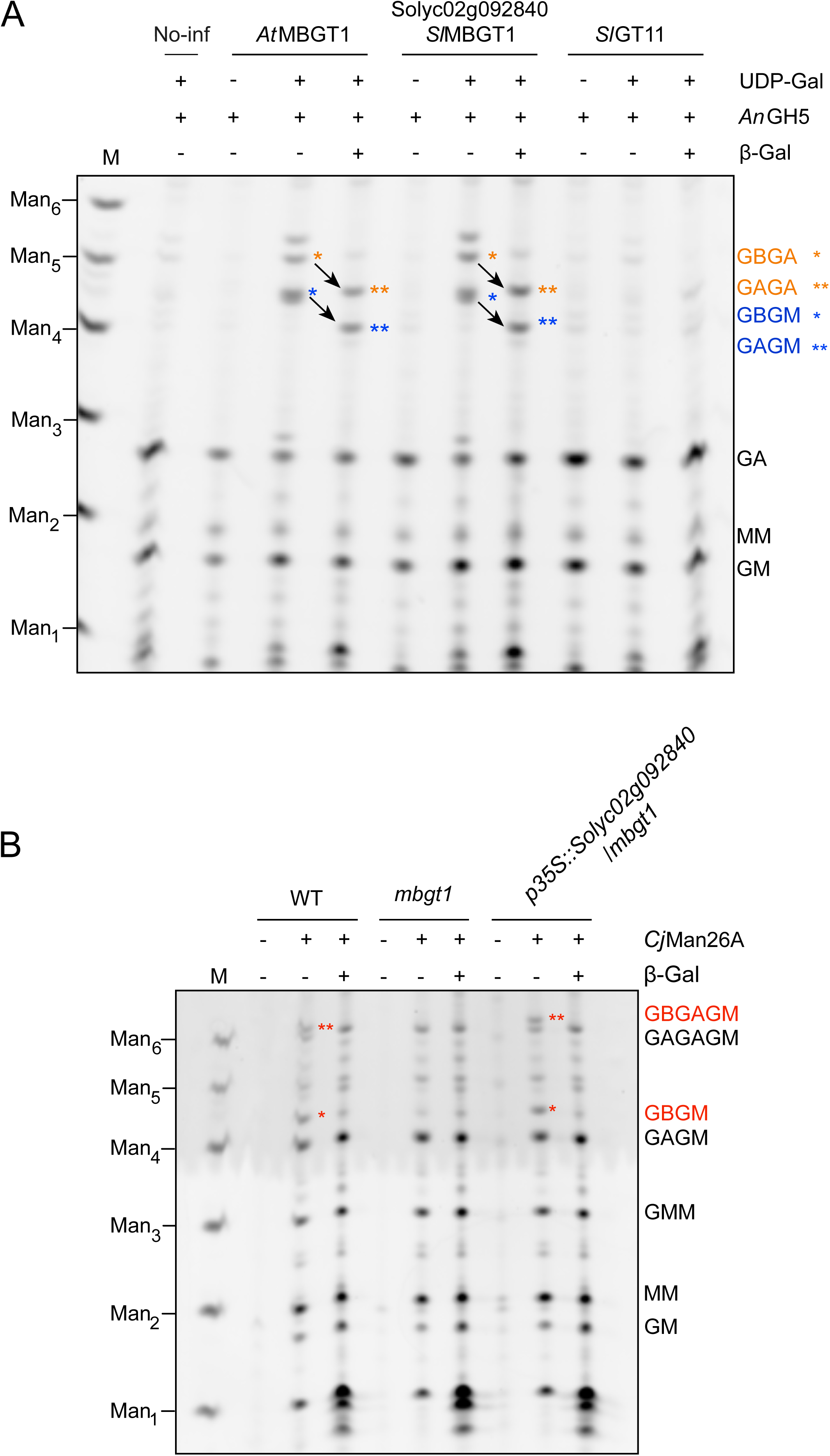
Biochemical activity of Solyc02g092840. (A) *In vitro* activity of recombinant Solyc02g092840. No-inf: solubilised microsomal fraction prepared from uninfiltrated tobacco leaves (negative control). After incubation with solubilised microsomes with expressed enzyme and UDP-Gal, substrates were digested with *An*GH5 mannanase and oligosaccharides visualised by PACE. (B) *In vivo* activity of Solyc02g092840 in the Arabidopsis *mbgt1* background. Cell wall material was digested with *Cj*Man26, β-galactosidase where indicated, and then the oligosaccharides were analysed by PACE. Three complementation lines were tested and a representative line is shown. M: marker lane (standard manno-oligosaccharides)

To confirm MBGT activity *in vivo* we next performed complementation analysis. Solyc02g092840 was expressed in the Arabidopsis *mbgt1* mutant under a strong constitutive promoter. The mannan structure in cell walls of young stem from WT (positive control), *mbgt1* (negative control), and complementation lines (*_pro_35S*∷*Solyc02g092840*/*mbgt1*) was analysed using *Cj*Man26A digestion. This mannanase released larger diagnostic β-GGM-derived oligos distinct from those from digestion of the AcGGM that is also present in the stem cell walls (e.g. MM, GMM, Yu *et al*., 2022). The β-Gal-containing oligosaccharides GBGAGM and GBGM were visible in WT and the complementation line but these β-galactosidase-sensitive stuctures were absent in *mbgt1* which produced only the α-galactosylated oligosaccharides GAGAGM and GAGM (Fig. 3B). Together with the *in vitro* activity evidence, we concluded that Solyc02g092840 encodes a MBGT that can contribute to the biosynthesis of β-GGM. Thus, we name it *Sl*MBGT1.

### Genes from asterids form a putative MBGT subclade in GT47A-III

The discovery that *Sl*MBGT1 in tomato (an asterid) is in the GT47A-III subclade implies that this enzyme emerged separately from *At*MBGT1 in Arabidopsis (a rosid) which resides in the GT47A-VII subclade. We considered whether the GT47A-III group of putative MBGT enzymes might be present more widely in plant lineages beyond tomato. Our previous paper reported asterid enzymes in three subclades within the GT47A-III defined as GT47A-III*a*, GT47A-III*b*, and GT47A-III*c* (Wilson *et al*., 2023). *Sl*MBGT1 is found in the previously uncharacterised subclade GT47A-IIIb. The xyloglucan galactosyltransferases such as AtXLT2 are associated with GT47A-IIIa. Xyloglucan arabinofuranosyltransferases from tomato (*Sl*XST1,2) and cranberry (*Vm*XST1) as well as xyloglucan β-xylosyltransferases from Robusta coffee (*Cc*XBT1) and blueberry (*Vc*XBT) are found in the *c* subclade (Immelmann *et al*., 2023; Wilson *et al*., 2023). Based on GT47 protein structural modelling and different activities found in these subclades, the putative donor specificity of GT47A-III subclades was speculated to vary with changes in five regions of amino acid sequence that form the donor sugar binding site (Wilson *et al*., 2023). Importantly, and in contrast to the XST and XBT sequences, the *Sl*MBGT-containing GT47A-IIIb subclade has greater similarity in these donor site residues to those of the characterised galactosyltransferases, consistent with the notion that the GT47A-IIIb clade consists of enzymes sharing the UDP-Gal donor specificity (Wilson *et al*., 2023). Genes from multiple orders of the asterid group (Ericales, Cornales, Lamiales, Solanales, Apiales, and Asterales) are found in this putative MBGT subclade, suggesting that other asterids may also have MBGT enzymes and have the capacity to synthesise β-GGM.

### Non-Solanales asterids exhibit β-GGM

As the phylogenetic analysis suggested the presence of MBGT enzymes in different orders of asterids, these plants should be capable of synthesising β-GGM. To test this, we collected tissues from blueberry (*Vaccinium corymbosum*, Ericales), carrot (*Daucus carota*, Apiales), olive (*Olea europaea*, Lamiales), and wild sunflower (*Helianthus tuberosus*, Asterales) and analysed their mannan structures (Fig. 4). Coinciding with our prediction, β-galactosylated β-GGM was found in blueberry and olive fruits. Since blueberry is within Ericales, an early branching order of asterids, this suggests the MBGT activity was already present early in asterid evolution. Interestingly, the subterranean parts from carrot and *H. tuberosus* had unsubstituted β-GGM, which was digested by the mannanase to GM and GMGM. Perhaps the β-GGM in these tissues is trimmed by galactosidases, as seen in Arabidopsis seed mucilage β-GGM (Yu et al., 2022). Consistent with the previous identification of β-GGM in kiwifruit (Schröder *et al*., 2001; Yu *et al*., 2022), a plant in the asterid Ericales group, this mannan structural analysis suggests that asterid orders possess β-GGM, and so have MBGTs, and therefore the other asterid enzymes in the GT47A-IIIb subclade may also have MBGT activity.

**Figure 4.**
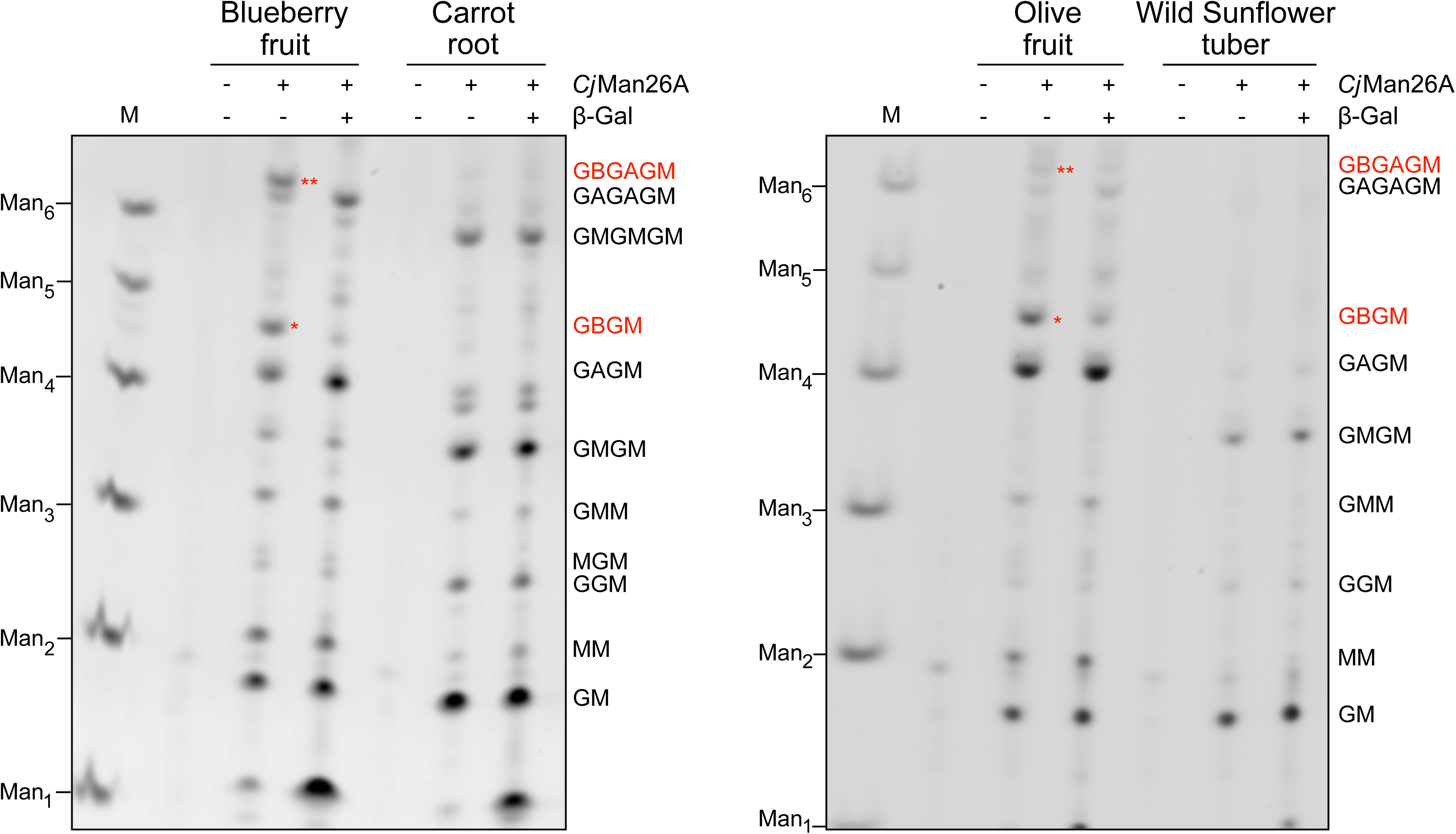
Mannan structure analysis of four species of asterids. Cell wall material was digested with mannanase *Cj*Man26 and β-galactosidase where indicated, and then the oligosaccharides were analysed by PACE. Blueberries and olive fruits contained β-Galactosylated β-GGM; carrot roots and Wild sunflower (*Helianthus tuberosus*) tubers contained unsubstituted β-GGM. M: marker lane (standard oligosaccharides).

### Functional convergence likely occurred with different amino acid substitutions

Given that MBGTs likely originated separately in asterids and rosids, we investigated whether the two distinct MBGT subclades (GT47A-VII in rosids and GT47A-IIIb in asterids) share the same evolutionary pathway to acquire MBGT activity. If they evolved through a common pathway, convergent amino acid substitutions should be observed at the same protein sites between the two MBGT subclades to allow β-GGM acceptor specificity to replace the xyloglucan acceptor specificity. To test this hypothesis, we performed an amino acid convergence analysis (Fukushima and Pollock, 2023). First, based on phylogenetic tree containing multiple asterid and rosid species, possible asterid and rosid orthogroups with MBGT activity were identified (Supplementary Data S1 Fig. 2). We do not know if there is redundancy in Arabidopsis AtMBGT1 paralogs (AT2G32750, AT2G32740, and AT2G31990), and no other rosid MBGTs have been identified, so it is not yet clear where MBGT activity diverged from xyloglucan galactosyltransferase activity presumably possessed by the common ancestor with GT47A-VII (Fig. 2) (Yu et al. 2022). We therefore considered several possible branch points between divergence from OsGT47A-VII and also more recent subclades. In asterids, the orthogroup containing *Sl*MBGT corresponds to the GT47A-IIIb subclade, and we therefore used this as the comparator for the convergence analysis. Convergence was examined between the stem branches of these orthogroups to understand which amino acid substitutions occured when MBGT activity most likely evolved (Supplementary Data 2). The results did not identify any clear convergence of amino acid sequence that could explain acquisition of the β-GGM acceptor specificity (Supplementary data S2). These findings suggest that while the two MBGT subclades are functionally convergent, their evolutionary pathways were not substantially shared. Thus, distinct amino acid substitutions in their stem branches, or changes that did not occur concurrently with the origins of the subclades, have likely contributed to the functional convergence of MBGTs.

## Discussion

### Recent acquisition of β-GGM

In this study, we identified a tomato MBGT located in GT47A-III, which is distinct from the subclade GT47A-III to which the rosid *At*MBGT1 belongs. This suggests the functional convergence of MBGT activity, and perhaps also convergence of β-GGM structure, during eudicot diversification. Furthermore, we found that asterid enzymes form a putative MBGT subclade (GT47A-IIIb) near the XST/XBT subclade (GT47A-IIIc). This subclade is asterid-specific (Wilson *et al*., 2023, Yu *et al*., 2022). Therefore, asterids diversified their GT47A-III enzymes after species divergence from other eudicot families. Interestingly, there are similarities in the evolutionary processes that led to MBGT activity in rosids and asterids. In asterids, GT47A-III is likely to have undergone three necessary gene duplication events. At the first divergence in a presumed clade of XLT2 xyloglucan galactosyltransferases, ancestors of XST/XBT/MBGT emerged. Then, the XST/XBT/MBGT ancestor split to create GT47A-IIIb (*Sl*MBGT1 subclade, β-GGM acceptor) and GT47A-IIIc (XST/XBT, xyloglucan acceptor). Interestingly, the rosid MBGT in GT47A-VII may also have evolved from a xyloglucan galactosyltransferase. *At*MBGT1 diverged from the common ancestor with *At*GT11 (GT47A-VII, suggested to be xyloglucan galactosyltransferase (Wei *et al*., 2021) and ultimately from a common ancestor with *At*MUR3 (GT47A-VI) xyloglucan galactosyltransferase (Fig. 2A). It therefore appears that the acquisition of MBGT activity each time involved a duplication event of genes encoding a xyloglucan galactosyltransferase, and subsequent accumulations of amino acid substitutions allowed the resulting β-galactosyltransferases to accept patterned galactoglucomannan in place of xyloglucan. Despite these similarities, we were unable to identify any convergent changes in amino acid sequence that might suggest critical residues for this change in acceptor binding.

### Do MBGTs from distinct subclades have different activity?

*At*XLT2 and *At*MUR3 are responsible for galactosylation of different positions on xyloglucan, where *At*XLT2 and *At*MUR3 add galactosyl residues on the second and third positions of a XXXG unit, respectively (Madson *et al*., 2003; Jensen *et al*., 2012). Given the two-fold screw conformation of the xyloglucan backbone (Levy, Maclachlan and Staehelin, 1997), the xylosyl residues of the XXXG unit position alternately on one side to the other of the backbone. In other words, two xylosyl residues of <u>X</u>X<u>X</u>G are on the opposite face to the one in the middle (X<u>X</u>XG), making the backbone appear xylosylated every other glucosyl residue on one side of the polymer. In contrast, the opposite side of the backbone appears xylosylated every four residues. It is therefore likely that *At*XLT2 and *At*MUR3 recognise the frequency of xylosyl residues on XyG and thus bind the xyloglucan substrate very differently to position the correct xylosyl residue for galactosylation. This notion led us to speculate that since the two MBGTs from rosids and asterids inherited their substrate recognition from the different ancestral xyloglucan galactosyltransferases, they might also bind their β-GGM substrates differently. In the case of β-GGM, the α-galactosyl substitutions of mannosyl residues always face just one side of the backbone due to the Glc-Man repeating unit of the backbone. When the mannosyl residues are frequently galactosylated (*e.g.*, GAGAGA), every other mannosyl residue of the backbone would be decorated, which structurally mirrors the side XyG backbone that *At*MUR3 recognises. Therefore, the MBGT enzymes that emerged in rosids (*e.g.*, *At*MBGT1) could be better able to recognise the frequently galactosylated β-GGM. On the other hand, *Sl*MBGT1, a descendant of XLT2 type galactosyltransferases, may favour the substrate decorated with less α-Gal on the β-GGM backbone such as GMGAGM patterns. Indeed, there seemed to be a difference in substitution patterns of β-GGM between Arabidopsis and tomatoes, where Arabidopsis showed a similar proportion of GBGA and GBGM while tomatoes primarily had GBGM with little GBGA (Yu *et al*., 2022). It would be interesting to investigate further whether the β-GGMs in and asterids have distinct substitution patterns and whether such structural differences change the functionality of β-GGM.

### Open question: What functions do glucomannans with different structures have?

We previously noted that β-GGM and xyloglucan have both structural and functional similaries (Yu *et al*., 2022). Here, we studied particularly the acquisition of the β-galactosylated disaccharide side chains in β-GGM, the synthesis of which now appears to have evolved at least twice in eudicots. Some structural variations of β-GGM have now been found: from the relatively highly β-galactosylated form in Arabidopsis primary cell wall (Yu *et al*., 2022) and blueberry fruit, the α-galactosylated β-GGM from Arabidopsis mucilage (Yu *et al*., 2018), and the unsubstituted form in carrot root and wild sunflower tuber (Fig. 4). These structural differences appear to be the result of biosynthetic gene expression variation, different specificity of MAGTs and MBGTs, and the production of hydrolases that trim the sidechain (*e.g.,* MUM2 β-galactosidase), rather than the absence of decoration enzymes in the genomes of these different eudicots. Although these structural variations may provide physiological adaptations, their specific role in growth and development remains to be elucidated. Despite the absence of some aspects of the structure in some organs, the convergent evolution of β-GGM biosynthesis to possess a similar structure in both rosids and asterids strongly suggests there has been an adaptive advantage to possession of the fully galactosylated β-GGM structure in eudicot cell walls. Why both xyloglucan and β-GGM have evolved this structure is an important outstanding question.

## Materials and Methods

### Plant materials

Arabidopsis and tobacco (*Nicotiana benthamiana*) were grown at 21°C under 16-hour day and 8-hour night conditions (Ishida *et al*., 2023). The Arabidopsis *mbgt1-1* mutant (SALK_065561) was as previously studied (Yu *et al*., 2022). Blueberry fruits and carrot roots were purchased at a local supermarket. Brined green olives were obtained from a supermarket and used after washing them with extensively with water. *Helianthus tuberosus* was grown in a home garden in Cambridge (UK) and subterranean parts were harvested in January 2023.

### Mannan structural analysis

Mannan structural analysis was conducted as described previously, (Ishida *et al*., 2023). Briefly, alcohol-insoluble residues and an alkali-soluble hemicellulose fraction were prepared. Then, mannan was digested sequentially by *An*GH5 or *Cj*Man26A mannanase and β-galactosidase. The digestion products were labelled with with 8-aminonaphthalene-1,3,6-trisulfonic acid and separated by carbohydrate gel electrophoresis.

### Molecular cloning and expression

Tomato gene expression data are from Tomato Genome Consortium (2012) accessed through the BAR expression browser https://bar.utoronto.ca/eplant_tomato/. For tobacco transient protein expression, the *SlMBGT1* gene was cloned into the pHREAC vector as previously described (structure: CaMV 35S promoter+CDS+eGFP+10xHis-tag+Nos terminator) (Peyret, Brown and Lomonossoff, 2019). *Sl*GT11 was cloned into the pEAQ vector (structure: CaMV 35S promoter+CDS+eGFP+6xHis-tag+Nos terminator) (Sainsbury, Thuenemann and Lomonossoff, 2009). The *SlMBGT1* expression vector for *in vivo* complementation was constructed by a Golden Gate-based system (Engler *et al*., 2009), in which the strategy and other cloning parts were the same as described previously (structure: CaMV 35S promoter +CDS+myc-tag+Nos terminator) (Ishida *et al*., 2023).The coding sequences of *Sl*GT11 and *Sl*MBGT1 were prepared by a commercial gene synthesis service (Integrated DNA Technologies, IA, USA). Nucleotide sequences of primers are described in Supplementary Data 3.

### In vitro assay of MBGT

Seven-week-old tobacco leaves were infiltrated with the Agrobacterium AGL1 strain harbouring the constructs described above. Microsome preparation and immunoblotting were performed as described previously (Ishida *et al*., 2023). For myc-tagged *At*MBGT1, Ab9106, a rabbit anti-Myc antibody (Abcam, Cambridge, UK), was used. For *Sl*MBGT1 and *Sl*GT11, Ab290 rabbit anti-GFP antibody (Abcam, Cambridge, UK) was used. Goat anti-rabbit IgG-HRP conjugates (Bio-Rad CA, USA) were used to detect primary antibodies. The *in vitro* assay using acceptor Arabidopsis mucilage β-GGM and donor UDP-Gal was carried out as described except that the reaction time was changed from 5 hours to overnight (Yu *et al*., 2022).

### In vivo complementation

Arabidopsis *mbgt1* (Yu *et al*., 2022) was transformed byAgrobacterium mediated floral dip using constructs to test MBGT activity as described previously (Ishida *et al*., 2023). Transformants were chosen by their seed eGFP signal and kanamycin resistance. Then, genotyping was carried out using primers in Supplementary Data 4.

### Phylogenetic analysis

The species phylogeny for 24 angiosperm species (ANA-grade: *Amborella trichopoda*, *Nymphaea colorata*, Monocots: *Oryza sativa*, Eudicots Rosids: *Arabidopsis thaliana* [Brassicales], *Brassica rapa* [Brassicales], *Capsella rubella* [Brassicales], *Citrus clementina* [Sapindales], *Eucalyptus grandis* [Myrtales], *Fragaria vesca* [Rosales], *Gossypium hirsutum* [Malvales], *Populus trichocarpa* [Malpighiales], *Theobroma cacao* [Malvales], *Vigna unguiculata* [Fabales], Eudicots Asterids: *Daucus carota* [Apiales], *Helianthus annuus* [Asterales], *Hydrangea quercifolia* [Rosales], *Lactuca sativa* [Asterales], *Lindenbergia philippensis* [Lamiales], *Olea europaea* [Lamiales], *Solanum lycopersicum* [Solanales], *Solanum tuberosum* [Solanales], *Vaccinium darrowii* [Ericales], *Vitis vinifera* [Vitales], other Eudicots: *Aquilegia coerulea* [Ranunculales]) with sequenced genomes was obtained from timetree.org. The phylogenetic analysis of GT47A-III was performed according to previous work (Fukushima and Pollock, 2023). In brief, genes homologous to the Arabidopsis GT47A-III were retrieved from the publicly available protein-coding gene sets of the 24 species using TBLASTX v2.9.0 with an E-value cutoff of 0.01 and >50% query coverage. In-frame codon sequence alignment was performed with MAFFT v7.455 (https://mafft.cbrc.jp/alignment/software/), ClipKIT v0.1.2 (https://github.com/JLSteenwyk/ClipKIT), and CDSKIT v0.9.1 (https://github.com/kfuku52/cdskit). The gene phylogeny was reconstructed using IQ-TREE v2.0.3 (https://github.com/iqtree/iqtree2) with the general time-reversible nucleotide substitution model and four gamma categories of among-site rate variation, and it was then reconciled with the species tree using GeneRax v1.2.2 (https://github.com/BenoitMorel/GeneRax).

### Amino acid convergence analysis

The reconciled gene tree and the codon alignment were used as inputs for CSUBST v1.1.0 (https://github.com/kfuku52/csubst) to infer the codon substitution history, which was then translated using the standard genetic code to obtain the probabilities of amino acid substitutions, as well as their convergence and divergence between focal branches. The inferred substitutions were mapped to the protein structure of *At*MBGT1 predicted by AlphaFold2 (Q84R16) using the ‘site’ function of CSUBST. Posterior probabilities of convergent or divergent substitutions greater than 0.5 were reported.

## Author Contributions

KI and PD conceptualised this study. KI and MP conducted biochemical experiments. KF conducted the amino acid convergence analysis. LFLW contributed to the screening of candidate *Sl*MBGT1 and phylogenetic analysis. YY and AEP supported the mutant generation. YY and LY provided resources and supported biochemical experiments. KI wrote the draft of the manuscript and PD, YY wrote a part of manuscript. All the authors revised the manuscript and approved the final version.

## References

Aspinall, G. O. (1959) “Structural Chemistry of the Hemicelluloses,” Advances in Carbohydrate Chemistry, pp. 429–468. doi: 10.1016/s0096-5332(08)60228-3.

Capek, P., Alföldi, J. and Lisková, D. (2002) “An acetylated galactoglucomannan from Picea abies L. Karst,” Carbohydrate research, 337(11), pp. 1033–1037.

Chernova, T. et al. (2020) “The Living Fossil Psilotum nudum Has Cortical Fibers With Mannan-Based Cell Wall Matrix,” Frontiers in plant science, 11, p. 488.

Engler, C. et al. (2009) “Golden gate shuffling: a one-pot DNA shuffling method based on type IIs restriction enzymes,” PloS one, 4(5), p. e5553.

Fernández, P. V. et al. (2012) “Sulfated β-d-mannan from green seaweed Codium vermilara,”Carbohydrate Polymers, pp. 916–919. doi: 10.1016/j.carbpol.2011.06.063.

Fukushima, K. and Pollock, D. D. (2023) “Detecting macroevolutionary genotype–phenotype associations using error-corrected rates of protein convergence,” Nature Ecology & Evolution. Nature Publishing Group, 7(1), pp. 155–170.

Gille, S. and Pauly, M. (2012) “O-Acetylation of Plant Cell Wall Polysaccharides,” Frontiers in Plant Science. doi: 10.3389/fpls.2012.00012.

Goubet, F. et al. (2003) “AtCSLA7, a cellulose synthase-like putative glycosyltransferase, is important for pollen tube growth and embryogenesis in Arabidopsis,” Plant physiology, 131(2), pp. 547–557.

Goubet, F. et al. (2009) “Cell wall glucomannan in Arabidopsis is synthesised by CSLA glycosyltransferases, and influences the progression of embryogenesis,” The Plant journal: for cell and molecular biology, 60(3), pp. 527–538.

Grieß-Osowski, A. and Voiniciuc, C. (2023) “Branched mannan and xyloglucan as a dynamic duo in plant cell walls,” Cell surface (Amsterdam). Elsevier BV, 9(100098), p. 100098.

Handford, M. G. et al. (2003) “Localisation and characterisation of cell wall mannan polysaccharides in Arabidopsis thaliana,” Planta, 218(1), pp. 27–36.

He, C. et al. (2017) “Cytochemical Localization of Polysaccharides in Dendrobium officinale and the Involvement of DoCSLA6 in the Synthesis of Mannan Polysaccharides,” Frontiers in plant science, 8, p. 173.

Immelmann, R., et al. (2023) “Identification of a xyloglucan beta-xylopyranosyltransferase from Vaccinium corymbosum,” Plant direct. Wiley, 7(7), p. e514.

Ishida, K. et al. (2023) “Differing structures of galactoglucomannan in eudicots and non-eudicot angiosperms,” PloS one, 18(12), p. e0289581.

Jensen, J. K. et al. (2012) “RNA-Seq Analysis of Developing Nasturtium Seeds (Tropaeolum majus): Identification and Characterization of an Additional Galactosyltransferase Involved in Xyloglucan Biosynthesis,” Molecular Plant, pp. 984–992. doi: 10.1093/mp/sss032.

Levy, S., Maclachlan, G. and Staehelin, L. A. (1997) “Xyloglucan sidechains modulate binding to cellulose during in vitro binding assays as predicted by conformational dynamics simulations,” The Plant journal: for cell and molecular biology, 11(3), pp. 373–386.

Liepman, A. H. et al. (2007) “Functional genomic analysis supports conservation of function among cellulose synthase-like a gene family members and suggests diverse roles of mannans in plants,” Plant physiology, 143(4), pp. 1881–1893.

Madson, M. et al. (2003) “The MUR3 gene of Arabidopsis encodes a xyloglucan galactosyltransferase that is evolutionarily related to animal exostosins,” The Plant cell, 15(7), pp. 1662–1670.

McCleary, B. V. et al. (1985) “The fine structures of carob and guar galactomannans,” Carbohydrate research, 139, pp. 237–260.

Meier, H. and Reid, J. S. G. (1982) “Reserve Polysaccharides Other Than Starch in Higher Plants,” in Loewus, F. A. and Tanner, W. (eds.) Plant Carbohydrates I: Intracellular Carbohydrates. Berlin, Heidelberg: Springer Berlin Heidelberg, pp. 418–471.

Moreira, L. R. S. and Filho, E. X. F. (2008) “An overview of mannan structure and mannan-degrading enzyme systems,” Applied microbiology and biotechnology, 79(2), pp. 165–178.

de O. Petkowicz, C. L., et al. (2001) “Linear mannan in the endosperm of Schizolobium amazonicum,” Carbohydrate polymers, 44(2), pp. 107–112.

Painter, T. J. (1983) “Structural evolution of glycans in algae,” Journal of Macromolecular Science, Part A:Pure and Applied Chemistry. De Gruyter, 55(4), pp. 677–694.

Peyret, H., Brown, J. K. M. and Lomonossoff, G. P. (2019) “Improving plant transient expression through the rational design of synthetic 5’ and 3’ untranslated regions,” Plant methods, 15, p. 108.

Preston, R. D. (1979) “Polysaccharide Conformation and Cell Wall Function,” Annual review of plant physiology. Annual Reviews, 30(1), pp. 55–78.

Sainsbury, F., Thuenemann, E. C. and Lomonossoff, G. P. (2009) “pEAQ: versatile expression vectors for easy and quick transient expression of heterologous proteins in plants,” Plant biotechnology journal, 7(7), pp. 682–693.

Scheller, H. V. and Ulvskov, P. (2010) “Hemicelluloses,” Annual review of plant biology, 61, pp. 263–289.

Schröder, R. et al. (2001) “Purification and characterisation of a galactoglucomannan from kiwifruit (Actinidia deliciosa),” Carbohydrate research, 331(3), pp. 291–306.

Schultink, A. et al. (2013) “The Identification of Two Arabinosyltransferases from Tomato Reveals Functional Equivalency of Xyloglucan Side Chain Substituents,” Plant Physiology, 163(1) pp. 86–94.

Terrett, O. M. et al. (2019) “Molecular architecture of softwood revealed by solid-state NMR,” Nature communications, 10(1), p. 4978.

Tomato Genome Consortium (2012) “The tomato genome sequence provides insights into fleshy fruit evolution” Nature 485 (7400): 635–641.

Voiniciuc, C. et al. (2015) “Highly Branched Xylan Made by IRREGULAR XYLEM14 and MUCILAGE-RELATED21 Links Mucilage to Arabidopsis Seeds,” Plant physiology, 169(4), pp. 2481–2495.

Voiniciuc, C. et al. (2016) “Extensive Natural Variation in Arabidopsis Seed Mucilage Structure,” Frontiers in plant science, 7, p. 803.

Voiniciuc, C. et al. (2019) “Mechanistic insights from plant heteromannan synthesis in yeast,” Proceedings of the National Academy of Sciences of the United States of America, 116(2), pp. 522–527.

Voiniciuc, C. (2022) “Modern mannan: a hemicellulose’s journey,” The New phytologist, 234(4), pp. 1175–1184.

Wang, Y. et al. (2013) “Identification of an additional protein involved in mannan biosynthesis,” The Plant journal: for cell and molecular biology, 73(1), pp. 105–117.

Wei, Q. et al. (2021) “The xyloglucan galactosylation modulates the cell wall stability of pollen tube,” Planta, 254(6), p. 133.

Wilson, L. F. L. et al. (2023) “The biosynthesis, degradation, and function of cell wall β-xylosylated xyloglucan mirrors that of arabinoxyloglucan,” The New phytologist. doi: 10.1111/nph.19305.

Ye, Z.-H. and Zhong, R. (2022) “Cell wall biology of the moss Physcomitrium patens,” Journal of experimental botany, 73(13), pp. 4440–4453.

Yu, L. et al. (2018) “The Patterned Structure of Galactoglucomannan Suggests It May Bind to Cellulose in Seed Mucilage,” Plant physiology, 178(3), pp. 1011–1026.

Yu, L. et al. (2022) “Eudicot primary cell wall glucomannan is related in synthesis, structure, and function to xyloglucan,” The Plant Cell. doi: 10.1093/plcell/koac238.

Zhong, R., Cui, D. and Ye, Z.-H. (2019) “Evolutionary origin of O-acetyltransferases responsible for glucomannan acetylation in land plants,” The New phytologist. Wiley, 224(1), pp. 466–479.

Zhu, L., Dama, M. and Pauly, M. (2018) “Identification of an arabinopyranosyltransferase from Physcomitrella patens involved in the synthesis of the hemicellulose xyloglucan,” Plant direct, 2(3), p. e00046.

