## Supplemental Files for "Convergent Acquisition of Glucomannan β-galactosyltransferases in Asterids and Rosids"

Supplementary data S1

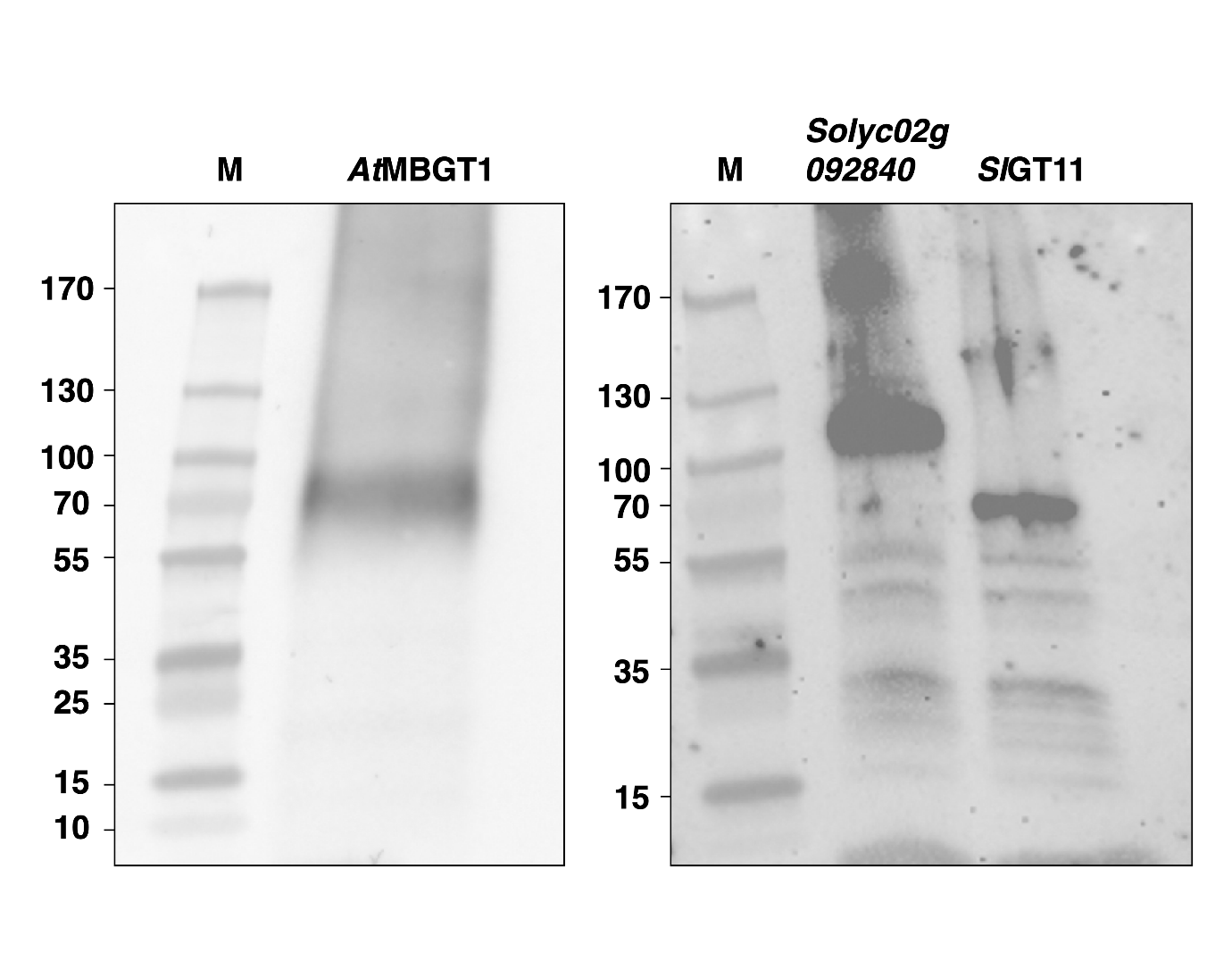

**Supplementary Data S1 Fig. 1.** Western blotting of the microsome fractions.

M: marker lane (standard proteins).

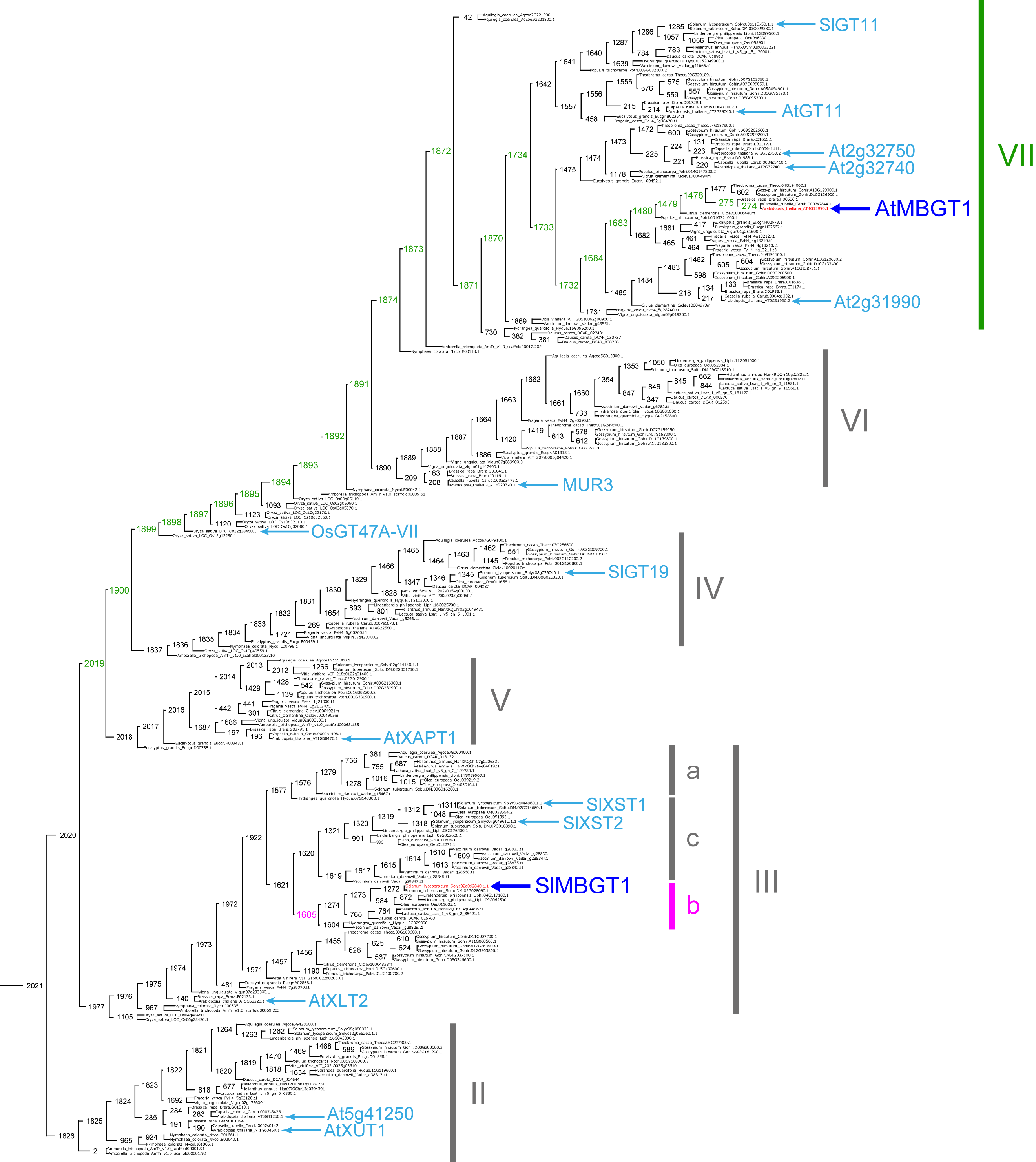

**Supplementary Data S1 Fig. 2.** The compared putative asterids MBGT subclade GT47A-IIIb and the putative rosids MBGT subclade branches are labelled in pink and green, respectively.

**Supplementary Data S1 Table 1.** Nucleotide sequences used in this study.

| No. | Name | Purpose | Sequence |
| --- | --- | --- | --- |
| 1 | SlMBGT | Protein expression | CACTCTGTGGTCTCAAATGATGCTGTCACCTTACTCCGACGAGTCGCCGGGCGCCGATGAAGCTATGCGCAAACCTCTAAAAACTAACAACAATGATTTTCTAAAGAATGCTGTGAGTAATTTTCAGTATCAGATTTCTACTCATCCTCGTTTCTGGCTTTTTACCTTCTTCCTTTTTTTTCAACTTGTGGTGCTTATCTTTACTCGCAATTCCCCTTTCCCTTTCTCAGTTCATTCCCCACCCAGTCCTCTCCCCCAATTCCCTTCAGAAATCGATGTATTTCCCGATTTACAGACAGCCCATCACGATGCAAATGTAATTTATCCCTTCGGCGATTCTGAATGCGAGTATGGTCGGGTCTATGTCTATAATTTACCATCCAAATTTAACAAAGATTTAGCTTTGTTGACCTGTGACGATCTAGACCCATGGAAGTGGCAGTGTGGTCTTGTGACCAATGATGGATATGGAAAGAGATCGACGGAGCTCGCCGGAATCTTGCCGGGGAATCTCTCCGCAGCGTGGTATCGTACAAATCAGTTCTCATCAGAGGTAATATTTCATTACCGGCTTTTAAACTATAGATGTAGAACCAACGACCCAGAATCTGCTACGGCTTTCTACATTCCGTTCTACGCCGGACAGGCTGTTGGGAAATACCTTTGGACTGATGAAATTGAAAATCGTGATTTTCTTAGCAACAAATTACTGAAATGGGTTCAAAGGCAAAAGTATTGGAAAAAATATAAGGGTTTGGACCATTTCCTCACACTTGGTCGGATAACGTGGGATTTCCGGCGATTAGGTGACCCAGAAAAACTCTGGGGATCCTCTTTTCTAAATCGACCGCAGATGCAGAATGTTACGCGATTCACAATCGAGAAGGCTCCATGGGATGCAAACGATATCAGTGTACCTTACCCTACCGGATTCCACCCTCATTCCGAGAAAGAACTCCGAGAATGGCAAAAGTTTGCACTTTCATATAACCGCACCAGTCTATTCACCTTCATAGGGGCGGCGCGTGGGGATATCGATAGTGATTTCAGATCGAGGTTAATGAGTTACTGTAGGAACGAGTCAGACTCGTGCCGAGTTGTGGACTGTGCCGTGATTCCCTGCTCTAACGGTTCATCAGAAATCCAAAAAGCACTTCTGAGCTCAGACTTCTGTTTACAGCCCAAAGGGGACAGCTTGACTCGGCGGTCAGTTTTTGATTGCATGGTGGCTGGTTCAGTACCGGTTTTCTTCTGGAGGCGAACGGCTTACACTCAGTACCAGTGGTTTTTGCCGGAGGACCCGGGGAGTTATTCCGTGTTTATAGACCCGGAAGCAGTGAGAAATGGGACAGCTTCCATTAAGGAAATTTTGATGAGTTACAGTAAAGATCAAGTAAGGAAAATGAGGGAAAAAGTTGTAGAGACAATTCCAAGAATAGTGTATGCAAGGCCAAGTGGAGGTCTTGGGAGTGTCAAGGATGCATTTGAAATTGCAATCGAAGGAGTATTGAAGAGGGTCAAGGACGAAAACGAATGGAAGGAATACGTGGATATGGGCAGTAGTTCGTGAGACCACGAAGTG |
| 2 | SlMBGT-F | Genotyping | TTCAGAAATCGATGTATTTCCCG |
| 3 | SlMBGT-R | Genotyping | ATTAACCTCGATCTGAAATCACTATCG |
| 4 | SlGT11 | Protein expression | CACTCTGTGGTCTCAAATGGATAACTCCCTCACAACAAGATCTCGCAATAAATTTTGGTTTGTTTTCTTTTTTCTATTTGTTTCTTGGTACTTATTGCTTTATGGAATTGATTGGTCATCTTTACCTGGTTTTGTAACAACGTCACGATATGAAGTAAATTCAATTGAATCCTTTCGTCCACCTCCTCATCATGAAAATGTTCATAATGTTAGCTCCAATTTCAATGCTACAAGTGTTGATGATGATAATGCTAGCAACGAGACGACGCGATCCAAGGAGGAAGAATCACTGTCTAATGAAAATGATAATGTGGTAGTACCTGATTTGGAGGAACTCCAGAAGGAATTGGAACCTTTGTTAAGAAAAATGGAACCTCCAAAAGAAGAGAAAAAGATCGAAAAGGAAGTTGAGAAGAAGGGAGGAAAATGTGCAGGACGATATATTTATGTGGCAGAAATACCTAGTAAGTTCAATGAAATCATGTTGAAGGAATGTAAATTGTTGAATAAATGGGAAGATATGTGTCAATATTTGGTAAATATGGGGCTTGGTCCTGATCTTGGAAACCCTCAAAGGATTTTCATGAACAAAGGTTGGTATACTACGAATCAATTTTCGTTAGAAGTTCTGTTTCATAACAGAATGAAACAGTACGATTGTCTAACGAATGATTCTTCAGTAGCATCAGCAGTTTTCGTTCCATACTATTCAGGGTTCGATGTTGCTAGGTACTTGTGGGATGATTTCAACACATCAATGAGGGATGCTGGTGCAATTGAGGTCGCAAAGTTTCTCAAGGAAAAACCTGAATGGAAAACAATGTGGGGAAGAGATCATTTCATGATTGCTGGTAGAATTACTTGGGATTTTAGGAGAGGTATTGAGGAAGATTCAGCTTGGGGGAACAAGTTAATGTTGTTACCTGAGGCAAAGAACATGACTATCTTAACAATCGAATCGAGTCCTTGGAACAGGAATGATTTCGCGATTCCATATCCAACCTATTTCCATCCTTCAAGTGACAGTGATGTAGTACAGTGGCAGAACAGGATGAGGAAGTTAAGGAGGAGGGTGTTGTTTTCATTCGCGGGGGCTCCACGTCCTCAACTTGAGGACTCAATAAGAAGTGAAATCATGGAACAATGCTCAGCCACGAGGCGTAAATGCAAGCTATTGGAGTGTAAAGACTTACACAACAAATGCAACAAGCCTGAACATGTAATGAGGCTGTTTCAAAGTTCAATTTTTTGCTTACAACCCTCAGGGGATTCGTTCACTAGGAGGTCTACTTTCGATTCAATTTTGGCTGGTTGTATCCCAGTGTTTTTCACTCCTGGATCCGCGTATGTCCAATATATATGGCACTTGCCAAAAGATTACACCAAATACTCAGTGCTCATACCGGAAGATGATGTTAGGAAAAAGAAAGTGAGCATCGAAAATGTACTATCTAAAATACCAAAATCACAAGTAGCAGCAATGAGAGAAGAAGTGATAAAGCTTATACCAAATGTAGTTTATGCAGATCCAAGAACAAGATTGGAGACAGTTAAAGATGCATTTGATTTGGCAGTGAAAGGGGTTCTTGAAAGAGTGGATGTAATAAGAAAAGAGATGAGACAGGGCAAATATTCAAGTATGATATTTGATGAAGAATTTAGCTGGAAGTATCATACATTTGGGACATTACAAAAGCATGAGTGGGATAGTTTCTTTTTGAGAACCAACAAAGAGAAGTACAGTTCGTGAGACCACGAAGTG |
| 5 | SlGT11-F | Genotyping | CTCTTTCAAGAACCCCTTTCACTGC |
| 6 | SlGT11-R | Genotyping | AGTGAAATCATGGAACAATGCTCAGC |
| 7 | pHREAC-SlMBGT-F | Protein expression | CACTCTGTGGTCTCAAAAATGATGCTG |
| 8 | pHREAC-SlMBGT-R | Protein expression | CACTTCGTGGTCTCACGAA |
